# Structural Basis for the Inhibition of the RNA-Dependent RNA Polymerase from SARS-CoV-2 by Remdesivir

**DOI:** 10.1101/2020.04.08.032763

**Authors:** Wanchao Yin, Chunyou Mao, Xiaodong Luan, Dan-Dan Shen, Qingya Shen, Haixia Su, Xiaoxi Wang, Fulai Zhou, Wenfeng Zhao, Minqi Gao, Shenghai Chang, Yuan-Chao Xie, Guanghui Tian, He-Wei Jiang, Sheng-Ce Tao, Jingshan Shen, Yi Jiang, Hualiang Jiang, Yechun Xu, Shuyang Zhang, Yan Zhang, H. Eric Xu

**Author notes:** Co-corresponding authors: H. Eric Xu,; or Yan Zhang; or Shuyang Zhang; or Yechun Xu. These authors contributed equally to this work (Wanchao Yin, Chunyou Mao, Xiaodong Luan, Dan-Dan Shen, Qingya Shen, Haixia Su).

## Abstract

The pandemic of Corona Virus Disease 2019 (COVID-19) caused by SARS-CoV-2 has become a global crisis. The replication of SARS-CoV-2 requires the viral RNA-dependent RNA polymerase (RdRp), a direct target of the antiviral drug, Remdesivir. Here we report the structure of the SARS-CoV-2 RdRp either in the apo form or in complex with a 50-base template-primer RNA and Remdesivir at a resolution range of 2.5-2.8 Å. The complex structure reveals that the partial double-stranded RNA template is inserted into the central channel of the RdRp where Remdesivir is incorporated into the first replicated base pair and terminates the chain elongation. Our structures provide critical insights into the working mechanism of viral RNA replication and a rational template for drug design to combat the viral infection.

---

The pandemic of COVID-19 caused by infection of SARS-CoV-2 has become a humanitarian crisis, with over 1.5 million infections and 87,000 death as reported on April 8^th^ of 2020(*1, 2*). SARS-CoV-2 is closely related to severe acute respiratory syndrome coronavirus (SARS-CoV) and several members from the beta coronavirus family, including bat and pangolin coronaviruses(*3-5*). Possibly due to the stronger binding affinity of the virus spike protein with the host receptor(*6-10*), SARS-CoV-2 appears to have much higher incidence of human to human transmission, resulting in much faster pace of infection worldwide. Tremendous challenges remain with ongoing battles against this notorious viral infection in many countries and finding an effective cure is an urgent need.

SARS-CoV-2 is a positive strand RNA virus, whose replication is mediated by a multi-subunit replication/transcription complex of viral non-structural proteins (nsp)(*11*). The core component of this complex is the catalytic subunit (nsp12) of an RNA-dependent RNA polymerase (RdRp)(*12, 13*). The nsp12 by itself has little activity and its functions require accessary factors including nsp7 and nsp8(*14, 15*), which increase the RdRp activity in template binding and processivity. The RdRp is also proposed to be the target of nucleotide class of antiviral drugs, including Remdesivir(*16-18*), which is a prodrug that is converted to the active drug in the triphosphate form (RTP) within cells(*19*). As such, RdRp has been a subject of intensive efforts of structural biology. The structures of nsp7, nsp8, and the complex of nsp12-nsp7-nsp8 have been determined(*15, 20-23*), providing an overall architecture of the RdRp complex assembly. However, no structure of the SARS-CoV-2 RdRp in complex with an RNA template or with nucleotide inhibitors has been solved, creating a knowledge gap in our understanding of working mechanism of RdRp and hampering the ongoing drug discovery effort against this viral infection. To fill in the above knowledge gap, we determined two cryo-EM structures of the SARS-CoV-2 RdRp complex either in the apo form or in a complex with a template-primer RNA and the antiviral drug Remdesivir.

For cryo-EM studies, we co-expressed nsp12 with nsp7 and nsp8 to form the core RdRp complex in insect cells (Figure 1A and Figure S1A-D). Stoichiometry ratio of nsp7 and nsp8 appeared to be less than nsp12, thus additional nsp7 and nsp8 from bacterial expression were supplemented before the final purification step to improve the yield of heterotrimeric complex. We also purified nsp12 alone and nsp12 by its own showed little activity in binding to a 50-base partial double-stranded template-primer RNA(Figure S1E), in agreement with previous studies with the SARS-CoV nsp12(*14*). The presence of nsp7 and nsp8 dramatically increased nsp12 activity to bind to the template-primer RNA (Figure S1E). The nsp12-nsp7-nsp8 complex also showed RNA polymerization activity on a poly-U template upon addition of adenosine triphosphate (ATP) in a time course-dependent manner (Figure 1B-1C). This RNA polymerization activity was effectively inhibited by the addition of the active triphosphate form of Remdesivir (RTP) (Figure 1D). Addition of 1 mM RTP in the presence of 10 mM ATP was able to completely inhibit the RdRp polymerization activity. Remdesivir, as a prodrug, at 5 mM concentration, did not have any inhibitory effect on the polymerization activity of the purified enzyme (Figure S1F).

**Figure 1.**
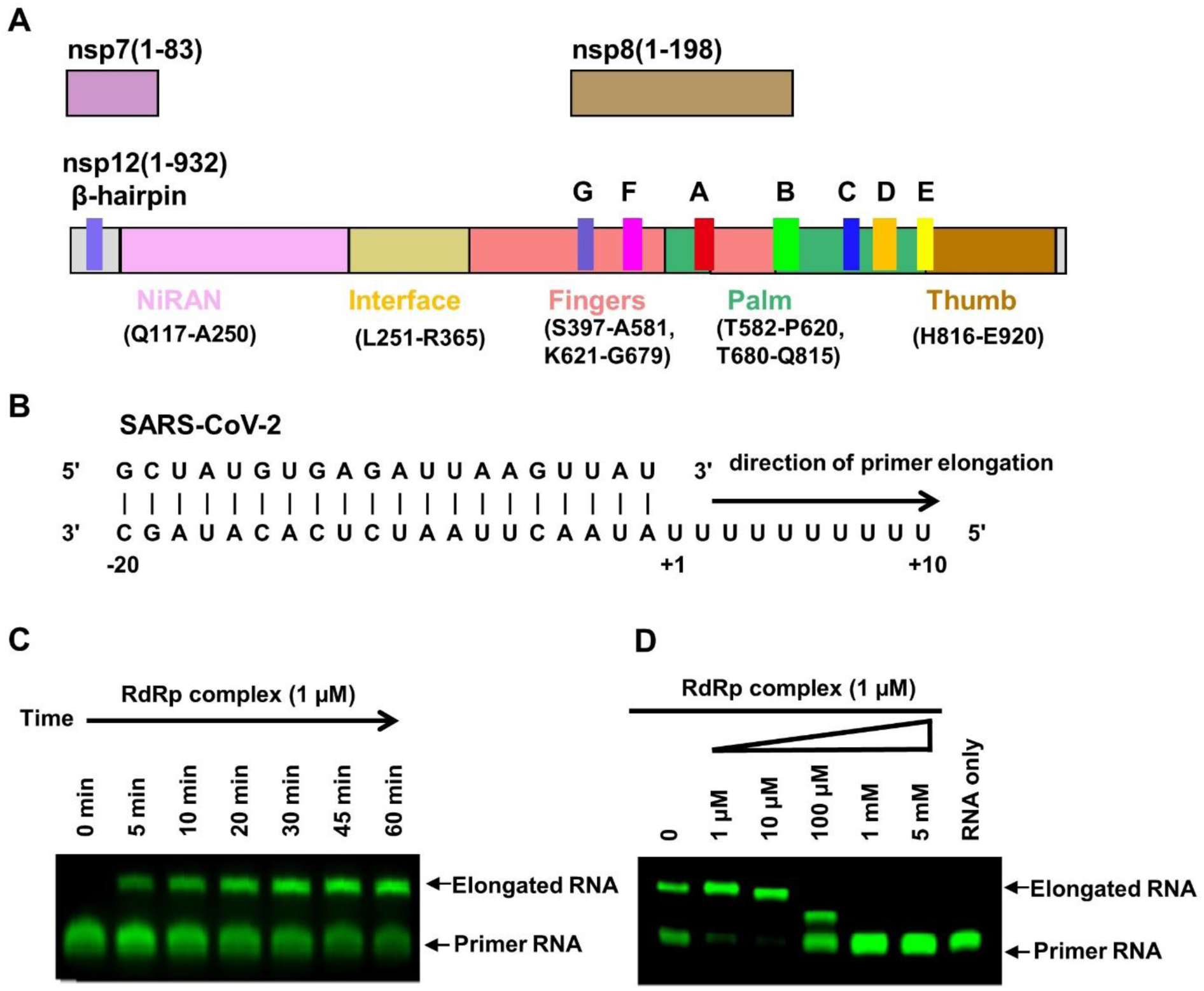
Assembly of an active nsp12-nsp-7-nsp8 RdRp complex and its inhibition by Remdesivir. A, Schematic diagram for the components of the active RdRp complex, containing nsp12, nsp7 and nsp8, with the nsp12 displayed in domain organization. The polymerase motif (A-G) and β-hairpin unique to the SARS-CoV-2 are highlighted. B, Sequence of RNA duplex with a 5’ U_10_ overhang as template used for extension reactions and RdRp-RNA complex assembly. C, The recombinant RdRp complex shows polymerase activity in vitro. The primer strand is labeled with fluorescence at the 5’ end. D, Elongation of partial RNA duplex by the purified RdRp complex is inhibited by RTP.

The purified RdRp complex is relatively thermostable with a melting temperature of 53°C (Figure S1G). Negative-stain EM visualization of the nsp12-nsp7-nsp8 complex displayed monodispersed particles in excellent homogeneity (Figure S1H). For the apo nsp12-nsp7-nsp8 complex, we vitrified the sample in the presence of the detergent DDM. The initial attempt of image processing reveals that the particles are preferentially oriented (Figure S2). Therefore, we collected and processed over 7400 micrograph movies of more than 5.7 million particle projections to increase the projection amount from the non-preferential orientation, out of which 81,494 particles are used to yield a density map of 2.8 Å resolution. Cryo-EM studies of the nsp12-nsp7-nsp8 complex bound to the template-primer RNA and RTP (termed as the template-RTP RdRp complex) were first hampered by the two facts. First, most particles were adsorbed to cryo-EM grid bars rather than staying in the vitreous ice. Second, RNA duplex dissociated from the template-RTP RdRp complex likely owning to the adverse effects during cryo-EM specimen preparation. Eventually, we prepared the cryo-EM specimen of the template-RTP RdRp complex at the concentration of 15 mg/ml supplemented with DDM prior to vitrification (Figure S3). We collected 2886 micrograph movies, which yielded a 2.5 Å resolution structure using 130,386 particle projections. Because of the relatively high resolution of our structures, EM density map is clear for key structure features across the complex (Figure S4).

The structure of the apo RdRp complex contains one nsp12, one nsp7 and two nsp8, with an overall arrangement resembling those seen in the SARS-CoV and the recently solved structure of SARS-CoV-2(*15, 23*) (Figure 2). Different from the SARS-CoV RdRp structure but similar to the recent SARS-CoV-2 RdRp structure, our structure reveals that nsp12 also contains an N-terminal β-hairpin (residues 31-50) and an extended nidovirus RdRp-associated nucleotidyl-transferase domain (NiRAN, residues 115-250)(*24*), with seven helices and a three β-strands (*15, 23*). Following the NiRAN domain is an interface domain (residues 251-365) composed of three helices and five β-strands, which is connected to the RdRp domain (residues 366-920). The nsp12 RdRp domain displays the canonical cupped right-hand configuration(*25*), with the finger subdomain (resides 397-581 and residues 621-679) forming a closed circle with the thumb subdomain (blue residues 819-920). The closed conformation is stabilized by the binding of nsp7 and nsp8, with one nsp8 molecule sitting on the top of the finger subdomain and interacting with the interface domain. The closed conformation of nsp12 is further stabilized by the nsp7-nsp8 heterodimer, which is packed against the thumb-index finger interface (Figure 2A-2B). In addition, we were able to assign two zinc ions in the conserved metal binding motifs composed by H295-C301-C306-C310 and C487-H642-C645-C646 (Figure 2C), which are also observed in the SARS-CoV RdRp structure(*15*), and these zinc ions likely to serve as conserved structural components in maintaining the integrity of RdRp architecture.

**Figure 2.**
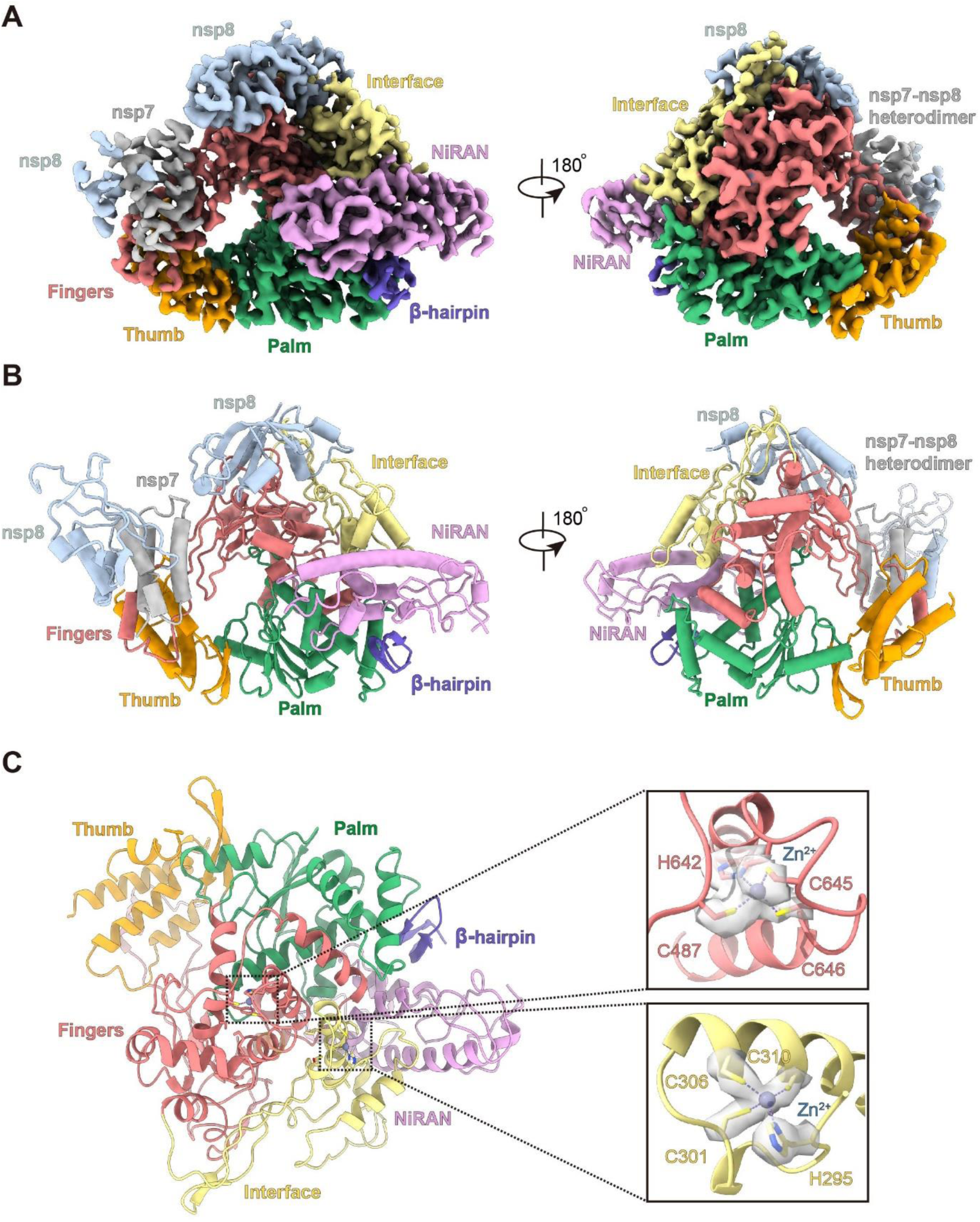
Cryo-EM Structure of the apo nsp12-nsp-7-nsp8 RdRp complex and structural features. A-B, Cryo-EM density map (A) and structure (B) of the nsp12-nsp7-nsp8 complex in apo state are shown in two views, colored according to Fig.1A. C, The conserved metal binding motifs composed by H295-C301-C306-C310 and C487-H642-C645-C646 are highlighted in the apo structure rendered in ribbon. To the right, the coordinate details of the zinc-binding residues are shown in stick against the cryo-EM map which is rendered in grey surface.

The structure of the template-RTP RdRp complex contains one nsp12, one nsp7 and one nsp8 (Figure 3). The second nsp8 in the apo structure appeared to be much more flexible in the template-RTP RdRp complex and it was not visible, therefore it was not included in the final model. In addition, the template-RTP RdRp structure contains 14 bases in the template strand, 11 bases in the primer strand, the inhibitor Remdesivir in its monophosphate form (RMP) (Figure 4), as well as a pyrophosphate and two magnesium ions that may serve as catalytic ions near the active site (Figure 5) (*26*).

**Figure 3.**
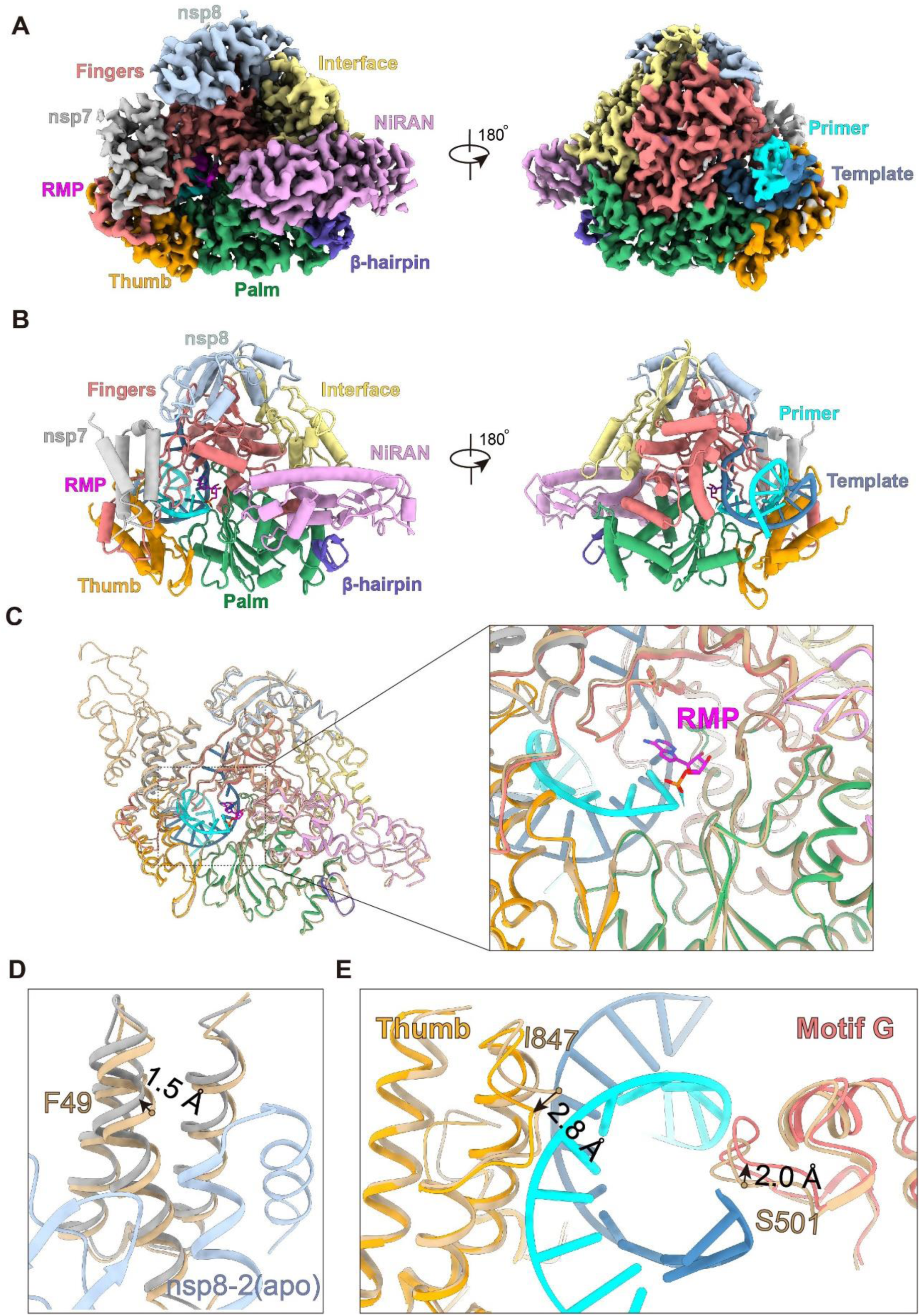
Cryo-EM Structure of the Remdesivir and template bound RdRp complex. A-B, Cryo-map (A) and model (B) are shown in two views for the nsp12-nsp7-nsp6 in complex with template RNA and Remdesivir. The nsp12-nsp-7-nsp8 trimer is colored as in Fig. 1A, Primer RNA in cyan and template RNA in dark blue. C, Superposition of the two complex structures is shown with the active site highlight in box. The apo state is shown in orange for clarity. D, Close view of the overlay of nsp7-nsp8 in two states with the nsp12 is omitted. E, Close view of the RNA exit tunnel in the apo and RNA-bound states.

**Figure 4.**
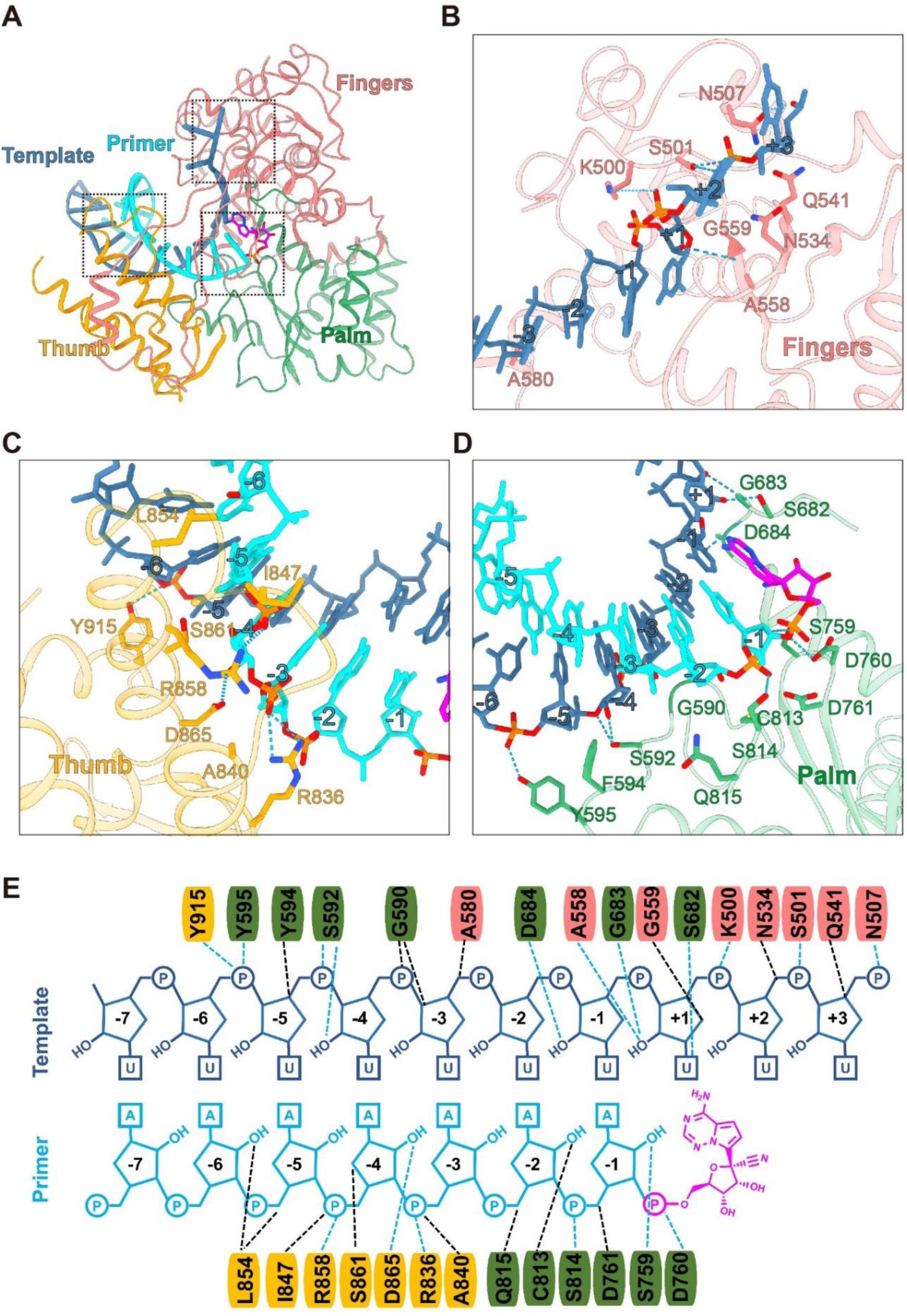
RNA Template Recognition by the RdRp complex. A, The interface between RNA template and RdRp. RNA template is shown as sticks against cryo-EM map. Detailed interactions between the RNA template and RdRp are shown in **B-D. E**, 2D representation of interactions between the template RNA and surrounding residues of RdRp.

**Figure. 5.**
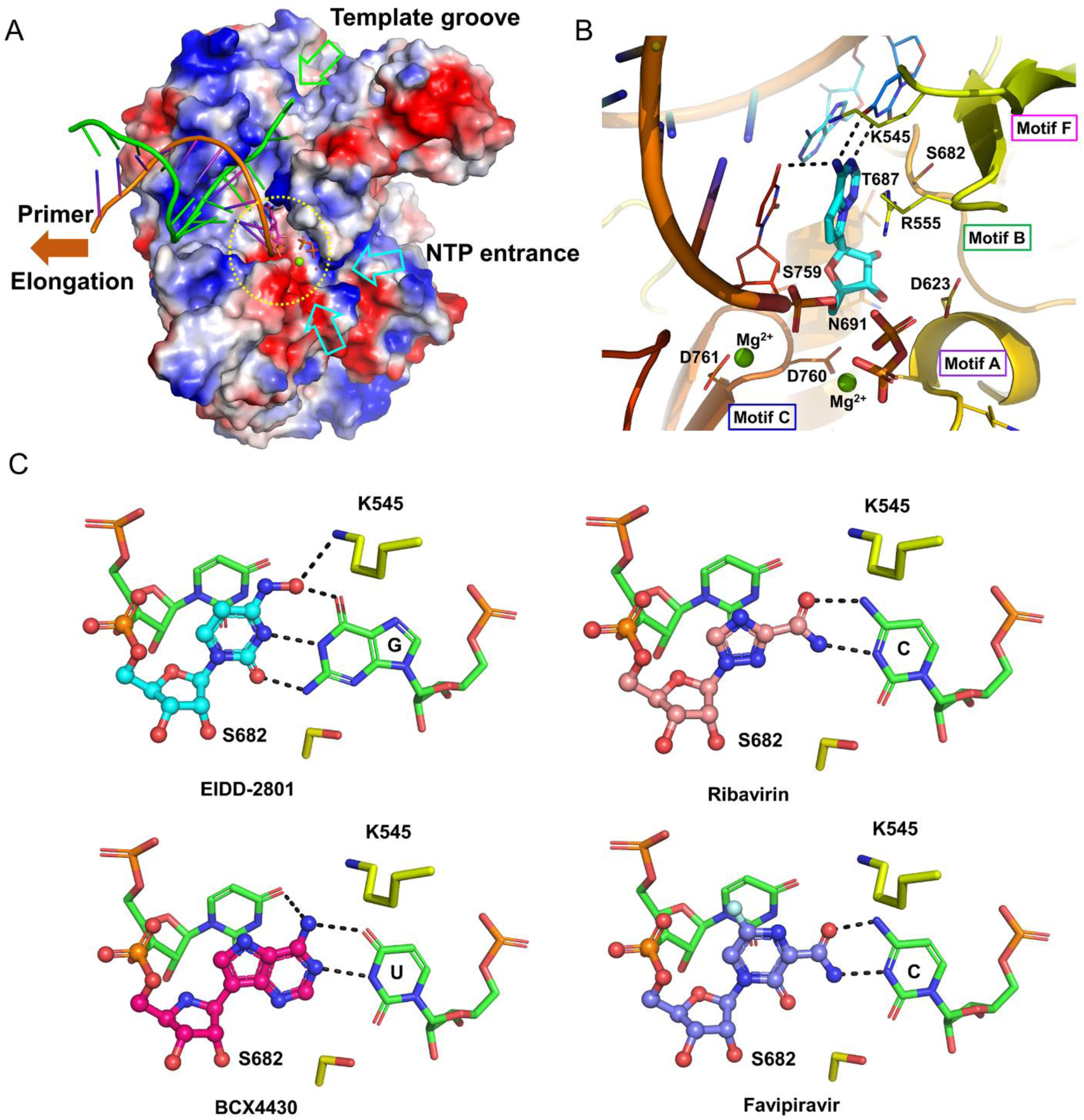
Structure of the RdRp active site and the binding modes of Remdesivir and other nucleotide analog antiviral drugs. A, Working mechanism of RdRp. Surfaces of the RdRp are colored according to their electrostatic potential from red (negative) to blue (positive). For clarity, residues 410-442, 834-919 of nsp12 and nsp8 are excluded from the figure. The template (green) primer (magenta) RNA are shown in cartoons. The covalently bound Remdesivir in a monophosphate form and the product, pyrophosphate, are shown as sticks. A magnesium ion and a water molecule at the active site are shown as a sphere and a ball, respectively. The active site is emphasized with a yellow dashed circle. The template groove, the entrance for NTP and the elongation direction are annotated with different colored arrows. B, The active site of RdRp covalently bound with the monophosphate form of Remdesivir, pyrophosphate and two magnesium ions. The protein and RNA are shown in cartoon representation, the covalently linked monophosphate form of Remdesivir and pyrophosphate are shown as sticks, two ions are shown as spheres, residues as well as bases interacting with Remdesivir are shown as thick lines. C, The binding modes of four nucleotide analogs, EIDD-2801, BCX4430, Ribavirin, and Favipiravir, with the RdRp, were generated based the complex structure of the RdRp with RTP. The nucleotide analogs are shown as balls and sticks, two residues together with two bases are shown as sticks.

The overall structure of the template-RTP RdRp complex is similar to the apo RdRp structure, with nsp12 in a closed conformation (Figure 2A and 3A). The double-stranded helix, formed by 11 base-pairs from the template-primer RNA (Figure 3B and Figure 4), is hold by the finger-palm-thumb subdomains. Extensive protein-RNA interactions are observed between the template-primer RNA and nsp12, with a total of 29 residues from nsp12 directly participating in the binding of the RNA (Figure 4E). Surprisingly no RNA interactions are mediated by nsp7 or nsp8 despite these two proteins are required for RNA binding by RdRp. Majority of protein-RNA interactions are mediated with the RNA phosphate-ribose backbones, with many interactions directly to 2’-OH groups (Figure 4E), thus providing a basis to distinguish RNA from DNA. There are no contacts from nsp12 to any base pairs of the template-primer RNA, suggesting no sequence-specific RNA binding by RdRp. This is consistent with the fact that no specific sequence is required for the enzymatic activity of RdRp at the elongation step.

At the 3’ end of the primer strand is Remdesivir monophosphate (RMP) (Figure 3C and 4D), which is covalently incorporated into the primer strand at the +1 position (Figure 4E). Additional nucleotides at +2 and +3 positions from the template strand from interactions with residues from the back from the index finger subdomain (Figure 4B). Despite the presence of excess amount of RTP in complex assembly, only a single RMP is assembled into the primer strand. Thus, Remdesivir, like many nucleotide analog prodrugs, inhibits the viral RdRp activity indirectly through non-obligate RNA chain termination, a mechanism that requires the conversion of the parent drug to the active triphosphate form(*27*).

The RMP position is at the center of the catalytic active site (Figure 5). The adenosine analog of RMP forms base-stacking interactions with upstream base from the primer strand and two hydrogen bonds with the uridine base from the template strand (Figure 5B and Figure S5). In addition, adenosine analog of RMP also forms interactions with side chains from K545 and R555. Near the bound RMP are two magnesium ions and a pyrophosphate. Both magnesium ions interact with phosphate diester backbone and they are part of catalytic active site. The pyrophosphate is at the gate of nucleotide entry channel to the active site and may block the entry of nucleotide triphosphate (NTP) to the active side (Figure 5A and 5B).

The catalytic active site of the nsp12 RdRp is constructed by seven conserved motifs from A to G (Figure 1 and Figure S6). Motifs ABCD are from the palm subdomain with the SDD sequence (residues 759-761) in motif C forming the catalytic active center (Figure 5B). Both D760 and D761 are involved in coordination of the two magnesium ions at the catalytic center. Motifs F and G are located within the finger subdomain, with motifs F and G interacting with the template strand RNA and directing this strand into the active site. Motif F also interacts with the primer strand RNA with its side chains from K545 and R555 contacting the base at the +1 position, thus stabilizing the incoming nucleotide in the correct position for catalysis. Compared with the structures of the poliovirus RdRp elongation complex(*28*) and the HCV Ns5b RdRp inhibitor complex(*29*), the mode of how template-primer RNA is orientated into the active site is exceedingly similar among these different viral RdRp complex (Figure S7). The residues involved in RNA binding as well as residues comprising the catalytic active site are highly conserved in this viral RdRp(*30, 31*), thus highlighting the highly conserved mechanism of genome replication by RdRp in these diverse RNA virus, and suggesting a possibility to develop broad spectrum antiviral inhibitors such as Remdesivir(*18*) and Galidisvir (BCX4430)(*32*).

Structural comparison reveals several interesting differences between the apo and complex structures (Figure 3D-3E). The first difference resides in the position of nsp7 appears to move toward the nsp core by 1.5 Å (as measured by nsp7 residue F49), leading to a rearrangement of the interface with the second nsp8, which results in the loss of the second nsp8 in the complex. The second difference is observed in the loop that connects the first and second helix of the thumb subdomain, which moves outward by as much as 2.8 Å (as measured nsp12 residue I847) to accommodate the binding of the double stranded RNA helix (Figure 3E). Motif G residues K500 and S501 also move outward by 2.0 Å to accommodate the binding of the template strand RNA. Outside of these changes, the apo nsp12 and the complexed nsp12 are exceedingly similar, with an RMSD of 0.52 Å for all Cα atoms across the whole protein. In particular, the structural elements that make up the catalytic active site can be superimposed identically (Figure S8), suggesting that SARS-CoV-2 RdRp is a relatively stable enzyme that is readily to function as a replicase upon the binding of RNA template. This is consistent with the fact that the purified RdRp complex had a relatively stable enzyme. Viral RdRp is a highly processive enzyme with rate of replication up to 100 nucleotides per second(*33*). No significant conformational changes between the apo and the active enzyme structures are consistent with the high processivity of the viral RNA polymerase, which does not need to consume additional energy for switch active site conformation during the replication cycle.

Besides Remdesivir, several drugs of nucleotide analogs, including Favipiravir, Ribavirin, Galidisvir, and EIDD-2801, have been shown to be efficient inhibitors in blocking SARS-CoV-2 replication in cell-based systems(*34, 35*). Like Remdesivir, these nucleotide analogs are proposed to inhibit viral RdRp indirectly through non-obligate RNA chain termination, a mechanism that requires the conversion of the parent compound to the triphosphate active form(*32*). The structure of the template-RTP RdRp complex provides an excellent model to rationalize how these drugs inhibit the SARS-CoV-2 RdRp activity (Figure 5C). In particular, EIDD-2801 has been shown to be 3-10 times more potent than Remdesivir in blocking SARS-CoV-2 replication(*35*). The N4 hydroxyl group off the cytidine ring forms an extra hydrogen bond with the side chain of K545. In addition, the cytidine base also forms an extra hydrogen bond with the guanine base from the template strand. These two extra hydrogen bonds may thus provide the explanation for the apparent higher potency for EIDD-2801 to inhibit SARS-CoV-2 replication.

The record-breaking infection by SARS-CoV-2 on a daily basis has inflicted emotional pain and economic burden across the globe. Enzymes that are vital for the viral life cycle are excellent antiviral drug targets as they are uniquely different from the host proteins. Among viral enzymes, RdRp is the major target of many existing nucleotide drugs. In this paper, we report the structure of the SARS-CoV-2 RdRp complex in the apo form and in the complex with a template-primer RNA and the active form of Remdesivir. The structures reveal how the template-primer RNA is recognized by the enzyme and the inhibition of the chain elongation by Remdesivir. Structure comparison and sequence alignment suggest that the mode of substrate RNA recognition and Remdesivir inhibition of RdRp are highly conserved in RdRp from diverse RNA viruses, providing a basis for designing broad spectrum antiviral drugs based on nucleotide analogs. Moreover, our structures also provide a solid template for modeling and modifying the existing nucleotide drugs, including the highly potent EIDD-2801. Together, these observations provide a rational basis to design even more potent inhibitors to combat the vicious infection of SARS-CoV-2.

## Materials and Methods

### Constructs and expression of the RdRp complex

The gene for the full-length SARS-CoV-2 nsp12 encompassing 932 residues was chemically synthesized with codon optimization (General Biosystems). The gene was cloned into a modified pFastBac baculovirus expression vector containing a 5’ ATG starting sequence and C-terminal Tobacco Etch Virus (TEV) protease site followed by a His8 tag. The plasmid construction contains one residue (methionine) to the N-terminus and GGSENLYFQGHHHHHHHH to the C-terminus of the protein. The genes for SARS-CoV-2 nsp7 encompassing the 83 residues and SARS-CoV-2 nsp8 encompassing the 198 residues were respectively cloned into the pFastBac vector containing a 5’ ATG starting sequence. All constructs were generated using the Phanta Max Super-Fidelity DNA Polymerase (Vazyme Biotech Co.,Ltd) and verified by DNA sequencing. All constructs were expressed in Spodoptera frugiperda (Sf9) cells. Cell cultures were grown in ESF 921 serum-free medium (Expression Systems) to a density of 2-3 million cells per ml and then infected with three separate baculoviruses at a ratio of 1:2:2 for nsp12, nsp7 and nsp8 at a multiplicity of infection (m.o.i.) of about 5. The cells were collected 48 h after infection at 27 °C and cell pellets were stored at − 80 °C until use.

In addition, the gene of nsp7 and nsp8 were respectively cloned into a modified pET-32a(+) vector containing a 5’ ATG starting sequence and C-terminal His8 tag for expression in E. coli. Plasmids were transformed into BL21(DE3)(Invitrogen). Bacterial cultures were grown to an OD600 of 0.6 at 37 °C, and then the expression was induced with a final concentration of 0.1 mM of isopropyl β-D-1-thiogalactopyranoside (IPTG) and the growth temperature was reduced to 16 °C for 18-20 h. The bacterial cultures were pelleted and stored at − 80 °C until use.

### Purification of the RdRp complex

For nsp7 or nsp8 expressed in bacterial, the purification was similar to the purification of nsp7 and nsp8 reported before(*15*). Briefly, the resuspended cells were lysed using a high-pressure homogenizer operating at 800 bar. Lysates were cleared by centrifugation at 25,000 × g for 30 min and were then bound to Ni-NTA beads (GE Healthcare). After extend washing with buffer containing 50 mM imidazole, the protein was eluted with buffer containing 300 mM imidazole. The eluted nsp7 sample was concentrated with a 3 kDa molecular weight cut-off Centrifuge Filter (Millipore Corporation) and then size-separated by a Superdex S75 10/300 GL column in 25 mM HEPES pH 7.4, 200 mM sodium chloride, 5% (v/v) glycerol. The eluted nsp8 sample was concentrated with a 30 kDa molecular weight cut-off Centrifuge Filter (Millipore Corporation) and then size-separated by a Superdex S75 10/300 GL column in 25 mM HEPES pH 7.4, 200 mM sodium chloride, 5% (v/v) glycerol. The fractions for the nsp7 or nsp8 were collected, concentrated to about 10 mg/ml, and stored at − 80 °C until use.

The insect cells containing the co-expressed RdRp complex were resuspended in Binding Buffer of 25 mM HEPES pH 7.4, 300 mM sodium chloride, 25 mM imidazole, 1 mM magnesium chloride, 0.1% (v/v) IGEPAL CA-630 (Anatrace), 1 mM tris(2-carboxyethyl)phosphine(TCEP) 10% (v/v) glycerol with additional EDTA-free Protease Inhibitor Cocktail (Bimake), and then incubated with agitation for 20 min at 4 °C. The incubated cells were lysed using a high-pressure homogenizer operating at 500 bar. The supernatant was isolated by centrifugation at 30,000×g for 30 min, followed by incubation with Ni-NTA beads (GE Healthcare) for 2 h at 4 °C. After binding, the beads were washed with 20 column volumes of Washing Buffer of 25 mM HEPES pH 7.4, 300 mM sodium chloride, 25 mM imidazole, 1 mM magnesium chloride, 1 mM TCEP and 10% (v/v) glycerol. The protein was eluted with 3-4 column volumes of Elution Buffer of 25 mM HEPES pH 7.4, 300 mM sodium chloride, 300 mM imidazole, 1 mM magnesium chloride, 1 mM TCEP and 10% (v/v) glycerol.

The eluted co-expressed RdRp complex was incubated with additional nsp7 and nsp8 from the bacterial expression in a 1:1:2 molar ratio and incubated at 4 °C for 4h. Incubated RdRp complex protein was concentrated with a 100 kDa molecular weight cut-off Centrifugal Filter (Millipore Corporation) and then size-separated by a Superdex S200 10/300 GL column in 25 mM HEPES pH 7.4, 300 mM sodium chloride, 0.1 mM magnesium chloride, 1 mM TCEP. The fractions for the monomeric complex were collected and concentrated for EM experiments of the apo complex. For the Remdesivir bound complex, the concentrated RdRp complex were diluted to 0.5 mg/ml with buffer of 25 mM HEPES pH 7.4, 100 mM sodium chloride, 2 mM magnesium chloride, 1 mM TCEP, combined with synthesized template-primer RNA and RTP in a 1:2:10 molar ratio and incubated at 4 °C for 2h. Then the incubated Remdesivir bound RdRp complex were concentrated for EM experiments.

### Cryo-EM data acquisition

Three microliters of the purified template and Remdesivir bound complex sample at about 15mg/ml (combined with 0.5 µL of 0.025% DDM) or the purified apo complex sample at about 0.75mg/ml (combined with 0.5 µL of 0.025% DDM) was applied onto a glow-discharged holey carbon grid (Quantifoil R1.2/1.3), blotted, and subsequently sample-coated grids were vitrified by plunging into liquid ethane using a Vitrobot Mark IV (Thermo Fischer Scientific). Cryo-EM imaging was performed on a Titan Krios equipped with a Gatan K2 Summit direct electron detector in the Center of Cryo-Electron Microscopy, Zhejiang University (Hangzhou, China). The microscope was operated at 300kV accelerating voltage, at a nominal magnification of 29,000× in counting mode, corresponding to a pixel size of 1.014 Å. 2,886 movies of the template and Remdesivir bound complex and 7,408 movies of the apo complex were obtained at a dose rate of about 8 electrons per Å^2^ per second with a defocus ranging from -0.8 to -3.0 μm. The total exposure time was set to 8s with intermediate frames recorded every 0.2s, resulting in an accumulated dose of 64 electrons per Å2 and a total of 40 frames per micrograph. The dose-fractionated image stacks were subjected to beam-induced motion correction using MotionCor2.1(*36*). A sum of all frames filtered according to the exposure dose in each image stack was used for further processing.

### Image processing and 3D reconstruction of the template and Remdesivir bound complex

Contrast transfer function parameters for each micrograph were determined by Gctf v1.06(*37*) and particle selection were performed using auto-picking command in RELION-3.0-beta2(*38*), yielding 2,438,446 particle projections. The resulted particles were imported to cryoSPARC v2.4.2(*39*) and the following reference-free 2D classification was performed producing 1,834,912 particle projections for further processing. This subset of particle projections was subjected to two rounds of Hetero Refinement and two selected subsets containing 580,293 projections were used to obtain a map with resolution of 2.9 Å in cryoSPARC. This portion of particle projections were imported back to RELION-3.0-beta2 and subjected to 3D refinement. Another round of 3D classification with mask on the complex produced a subset of 130,386 particle projections for the final reconstruction. After consecutive rounds of 3D refinement and Bayesian polishing of 130,386 particle projections, the final map has an indicated global resolution of 2.5 Å at a Fourier shell correlation (FSC) of 0.143. Local resolution was determined using the Bsoft package with half maps as input maps(*40*).

### Image processing and 3D reconstruction of the apo complex

Contrast transfer function parameters determination, particle selection, 2D and 3D classification were performed as above described. Briefly, auto-picking yielded 5,791,192 particle projections. The following reference-free 2D classification in cryoSPARC was performed and produced 2,160,423 particle projections for two rounds of Hetero Refinement. One selected subset containing 257,941 projections was used to obtain a map with resolution of 3.2 Å. The selected subset of particle projections was subsequently imported back to RELION-3.0-beta2 and subjected to consecutive rounds of 3D refinement and Bayesian polishing. Another round of 3D classification with mask on the complex produced a subset of 81,494 particle projections for the final reconstruction. The final map has an indicated global resolution of 2.8Å at a Fourier shell correlation (FSC) of 0.143.

### Model building and refinement

The model of the SARS-CoV RdRp (PDB code: 6NUR) was used as the start for model rebuilding against SARS-CoV-2 RdRp apo structure and RdRp-RNA-Remdesivir complex structure. The model was docked in the using Chimera(*41*), followed by iterative manual adjustment in COOT(*42*), real space refinement using real_space_refine in PHENIX(*43*). The model statistics was validated using MolProbity. Structural figures were prepared in Chimera and PyMOL (https://pymol.org/2/). The final refinement statistics are provided in Table S1.

### Thermal shift assay (TSA)

The thermal stability of the RdRp complex was evaluated by using TSA. Briefly, a mixture of 0.5 mg/ml protein and 10× SYPRO Orange (Thermo Fisher Scientific) was incubated for 10 min at room temperature. The reaction was performed in 96-well plates with a final volume of 20 μl.The thermal melting curve were monitored using a LightCycler 480 II Real-Time PCR System (Roche Diagnostics, Rotkreuz, Switzerland) with a ramp rate of 1 °C at the temperature range from 25 °C to 80 °C. The melting peak were calculated by the LightCycler 480 software provided by Roche Diagnostics.

### Gel mobility shift assay to detect RNA–RdRp protein binding

A gel mobility shift assay was performed to detect RNA binding by the RdRp complex. The binding reaction contained 25 mM HEPES pH 7.4, 100 mM sodium chloride, 2 mM magnesium chloride and 1 mM TCEP, 9 μg RdRp complex protein with increasing amounts of template-primer RNA (0, 0.3, 0.6, 1.2 and 2 μg). For the nsp12 protein, the binding reaction was combined with 1.5 μg template-primer RNA. Binding reactions were incubated for 30 min at room temperature and resolved on 4-20% native polyacrylamide gel (Thermo Fisher Scientific) running in 1×TBE buffer at 90 V for 1h in 4 °C cool-room. Then the gel was carefully taken out and was dyed with Ultra GelRed Nucleic Acid Stain (Vazyme Biotech Co.,Ltd) according to the manufacturer’s protocol. The dye-stsined gel was visualized on a BIO-RAD Fluorescence Imager.

### Preparation of template-primer RNA for polymerase assays

A short RNA oligonucleotide of sequence 5’-FAM-GCUAUGUGAGAUUAAGAAUU-3’ was used as the primer strand and a longer RNA oligonucleotide of sequence 5’-UUUUUUUUUUAAUUCUUAAUCUCACAUAGC -3’ was used as template strand. To anneal the RNA duplex, both oligonucleotides were mixed at equimolar ratios in annealing buffer (10 mM Tris-HCl pH 8.0, 25 mM NaCl and 2.5 mM EDTA), denatured by heating to 94°C for 5 min and allowed to slowly cool to room temperature.

### RdRp enzymatic activity assay and its inhibition by Remdesivir-triphosphate and Remdesivir

The purified SARS-CoV-2 RdRp complex from insect cell at final concentration of 1 μM was incubated with 3.0 μM dsRNA and 10 mM ATP in the presence of 1.14 U/μl RNase inhibitor in Reaction buffer containing 20 mM Tris pH8.0, 10 mM KCl, 6 mM MgCl2, 0.01% Triton-X100, 1 mM DTT, which were prepared with DEPC-treated water.The total reaction volume was 20 μl. After incubation for 0 min, 5 min, 10 min, 20 min, 30 min, 45 min, and 60 min at 37°C water bath, 40 μl quench buffer (94% formamide, 30 mM EDTA, prepare with 2x Reaction buffer) was added to stop the reaction. A sample of 18 μl of reaction was mixed with 2 μl 10x DNA loading buffer and half of the sample (10 μl) was loaded onto a 20% Urea-PAGE denatured gel and run at 120V for 1h. Image via Tanon-5200 Multi Fluorescence Imager.

The setup for the inhibition assays of the RdRp by RTP and Remdesivir is identical to the above for the RdRp enzymatic assays, except that RTP or Remdesivir were added to final concentration of 0 μM, 1 μM, 10 μM, 100 μM, 1 mM, and 5 mM for 60 min before the addition of 10 mM ATP.

## Supporting information

PDB for the apo structure

PDB for the Remdesivir structure

Map for the apo structure

Map for the Remdesivir bound structure

## Acknowledgments

The cryo-EM data were collected at the Center of Cryo-Electron Microscopy, Zhejiang University. This work was partially supported by Shanghai Municipal Science and Technology Major Project 2019SHZDZX02 and XDB08020303 to H.E.X.; Zhejiang University special scientific research fund for COVID-19 prevention and control E33 and the National Science Foundation of China 81922071 to Y.Z.; Science and Technology Commission of Shanghai Municipality 20431900100 and Jack Ma Foundation 2020-CMKYGG-05 to H.J. and J.S.; CAMS Innovation Fund for “13th Five-Year” National Science and Technology Major Project for New Drugs 2019ZX09734001-002 and Tsinghua University-Peking University Center for Life Sciences 045-160321001 to S. Z.; National Key Research and Development Program of China Grant 2016YFA0500600 and National Natural Science Foundation of China 31970130 and 3167083 to S.T.; and the National Key R&D Program of China 2016YFA050230 to Y.X; the National Natural Science Foundation 31770796 and the National Science and Technology Major Project 2018ZX09711002 to Y.J. We also thank MedChemExpress for making Remdesivir.

## Author contributions

W.Y. designed the expression constructs, purified the RdRp complex, prepared samples for negative stain and data collection toward the structures, and participated in figure and manuscript preparation. C.M. and D.-D.S. evaluated the specimen by negative-stain EM, screened the cryo-EM conditions, prepared the cryo-EM grids and collected cryo-EM images with the help of S.C.; D.-D.S. and C.M. performed density map calculations; Q.S. and H.S. participated in the model building and refined the final models; X.L. designed RdRp activity assays and Remdesivir inhibition experiments as well as expression constructs of the RdRp complex; X. W., F. Z., and M.G. participated in expression, purification and functional assays of the RdRp; Y-C.X, G. T., and J. S. made Remdesivir triphosphate form; Y.J. participated in experimental design and manuscript editing; H.J. conceived and coordinated the project; S.Z. conceived the project, initiated collaboration with H.E.X., and supervised X.L.; Y.X. analyzed the structure and modeling and participated in figure preparation; Y.Z. supervised Q.S., C.M., and D.D.S, analyzed the structures and participated in manuscript writing; H.E.X. conceived and supervised the project, analyzed the structures, and wrote the manuscript with inputs from all authors.

## Data and materials availability

Density maps and structure coordinates have been deposited with immediate release. The accession numbers of Electron Microscopy Database and the Protein Data Bank are EMD-30209 and PDB ID 7BV1 for the apo RdRp complex; EMD-30210 and PDB ID 7BV2 for the template RNA and Remdesivir bound RdRp complex. These files are immediately available for download as supplemental materials in this article.

## Supplemental Figures

**Figure S1.**
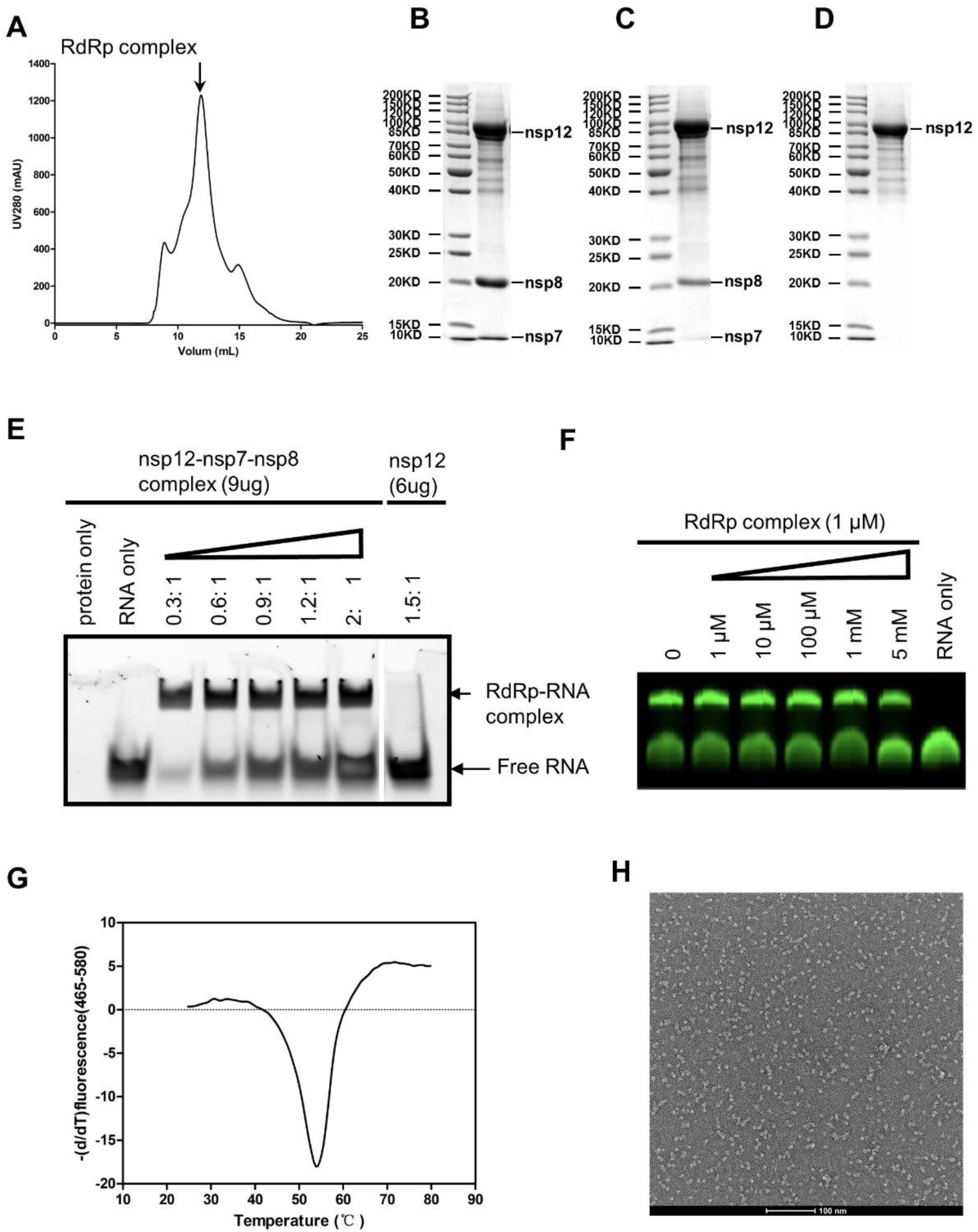
Purification and characterization of the RdRp complex. A, Gel filtration profile of the RdRp complex with additional nsp7 and nsp8, showing a sharp peak. B, SDS gel of the purified RdRp complex with additional nsp7 and nsp8, showing balanced ratios for each subunit. C, SDS gel of the co-expressed RdRp complex, showing under stoichiometry of nsp7 and nsp8. D, SDS gel of the purified nsp12. E, Gel mobility shift of the RdRp-RNA complex. F, Remdesivir in the prodrug form did not inhibit the activity of the purified RdRp complex. G, Thermal stability shift analysis of the RdRp complex, indicating its melting temperature of 53°C. H, Negative stain EM of the RdRp complex.

**Figure S2.**
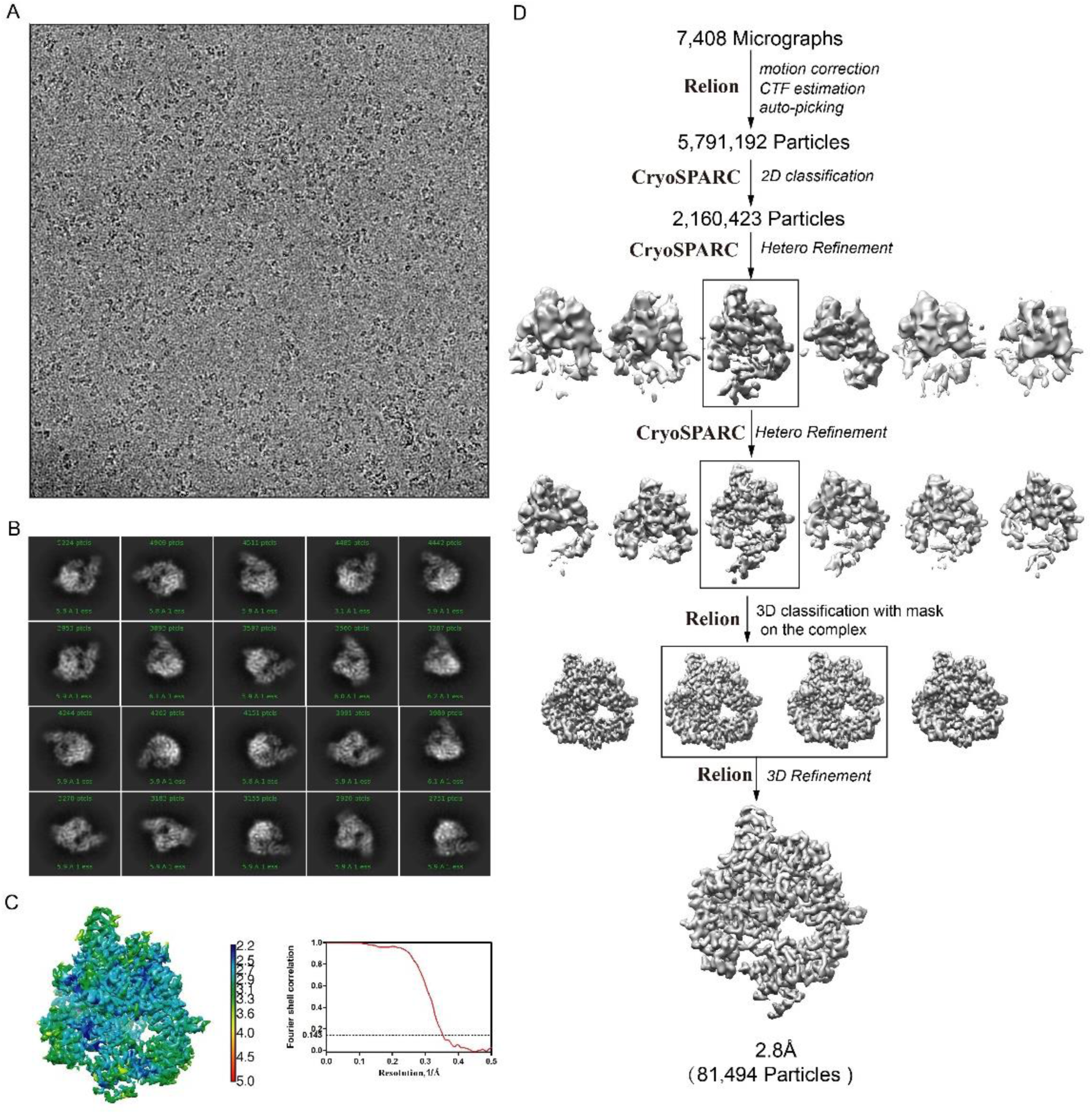
Single particle cryo-EM analysis of the apo complex. A, Representative cryo-EM micrograph of the apo complex. B, Representative 2D class averages of the apo complex. C, Cryo-EM map of the apo complex, colored by local resolution (Å) and ‘Gold-standard’ Fourier shell correlation curve. D, Flowchart of cryo-EM works of the apo complex.

**Figure S3.**
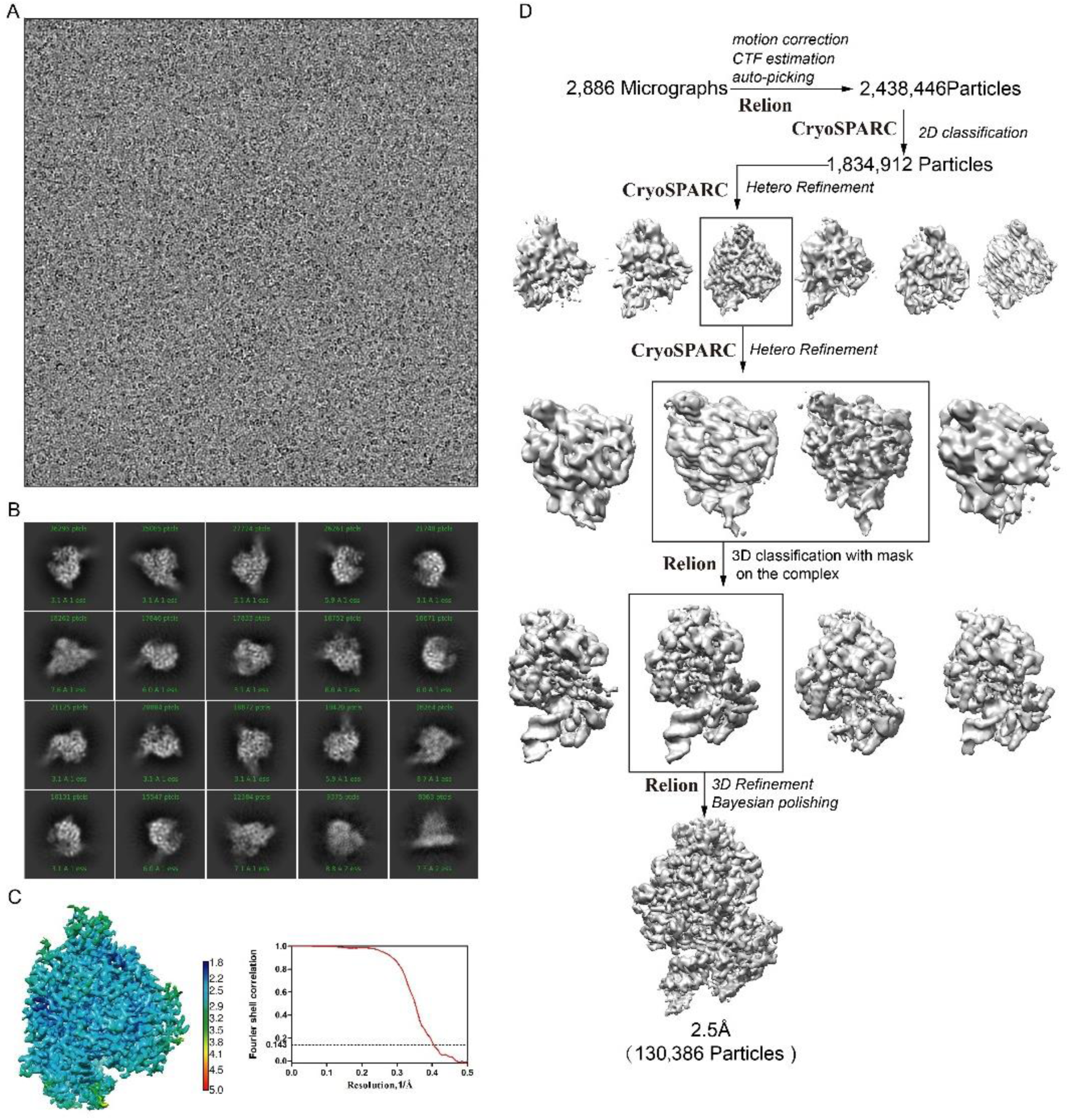
Single particle cryo-EM analysis of the template-RTP RdRp complex. A, Representative cryo-EM micrograph of the template-RTP RdRp complex. B, Representative 2D class averages of the template-RTP RdRp complex. C, Cryo-EM map of the template-RTP RdRp complex, colored by local resolution (Å) and ‘Gold-standard’ Fourier shell correlation curve. D, Flowchart of cryo-EM works of the template-RTP RdRp complex.

**Figure S4.**
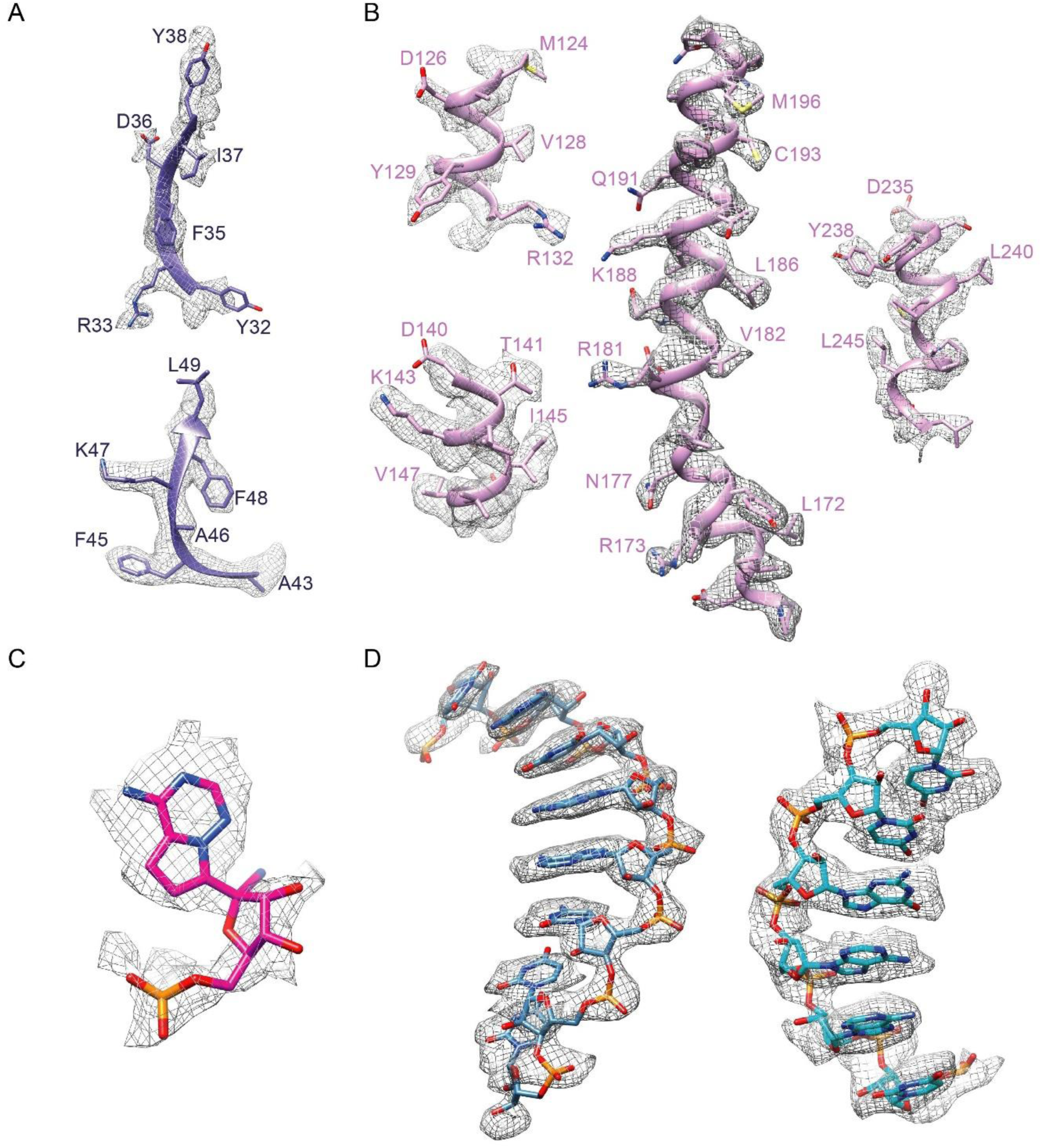
Quality of cryo-EM map for the RdRp in complex with RNA template and RTP. A, Cryo-EM density map and model of β-hairpin. B, Cryo-EM density map and model of NiRAN domain. C, Cryo-EM density map of Remdesivir. D, Cryo-EM density map and model of the template strand RNA and the primer strand RNA.

**Figure S5.**
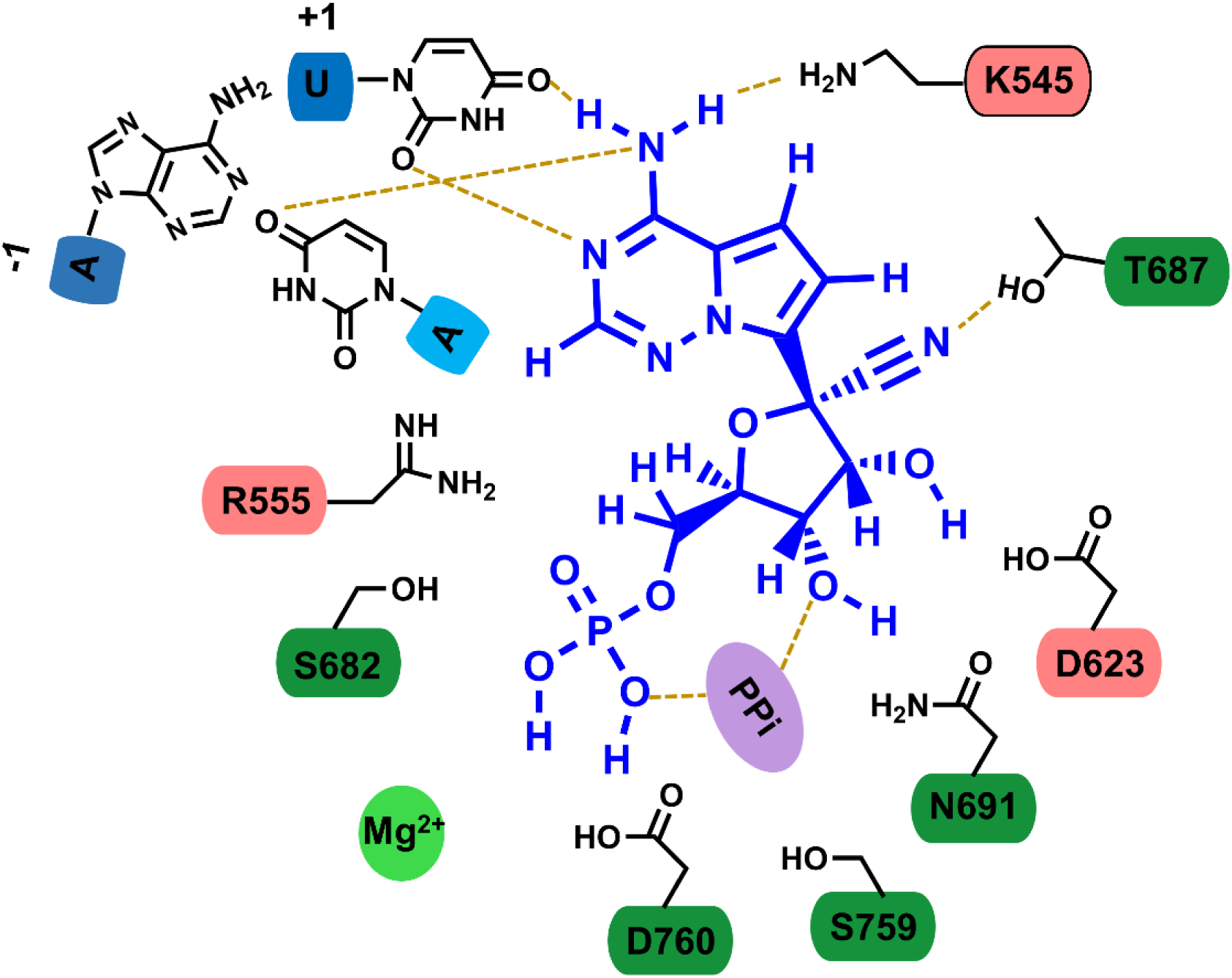
2D representation of interactions of the bound Remdesivir in a monophosphate form with surrounding residues and nucleotide bases.

**Figure S6.**
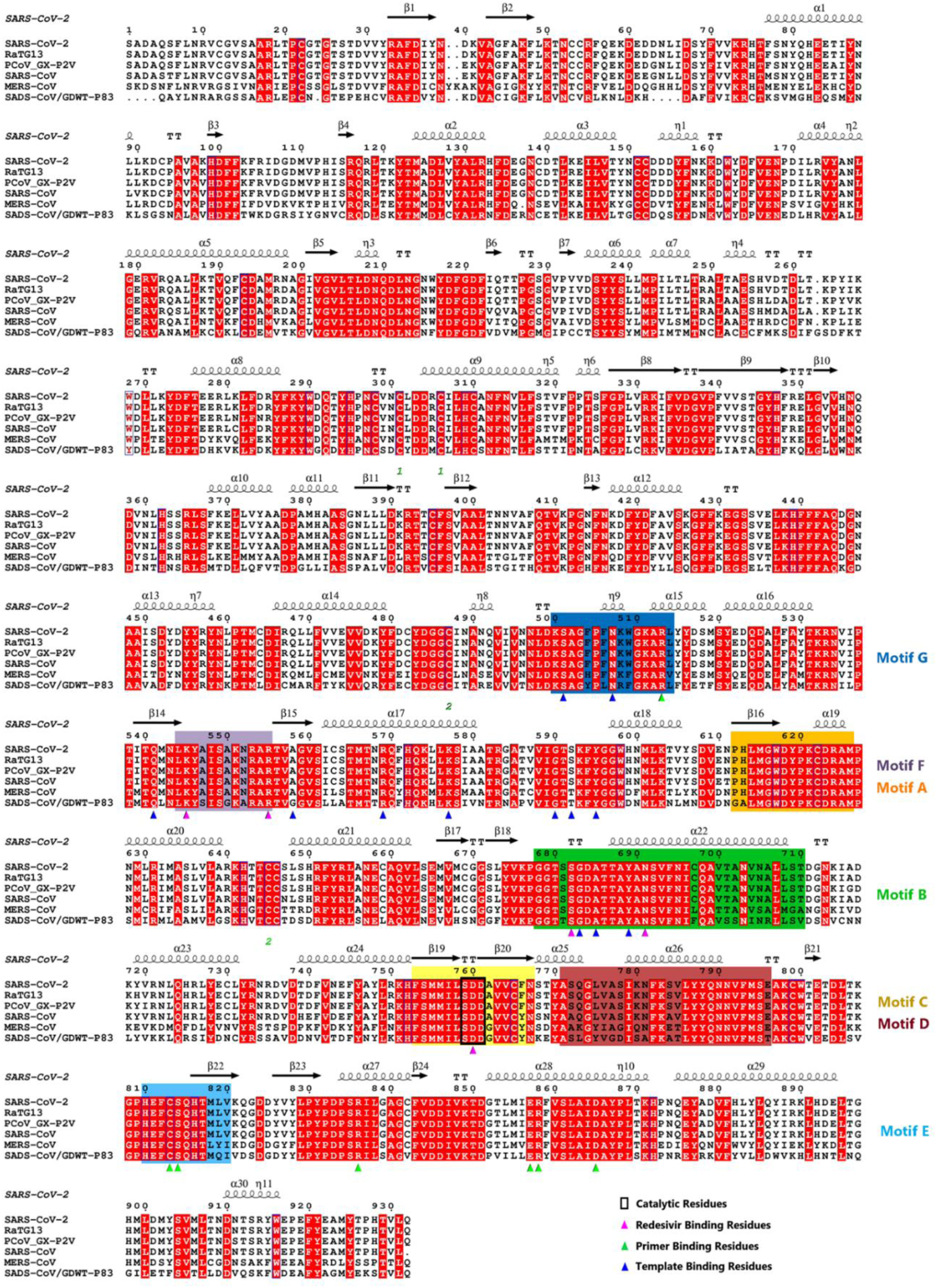
Sequence alignment of six coronavirus nsp12. Sequences of SARS-CoV-2, RaTG13, PCoV_GX-P2V, SARS-CoV, MERS-CoV, and SADS-CoV nsp12 were aligned with clustalw and ESPript. Secondary structures, seven conserved motifs, three catalytic residues, and residues interacting with the covalently linked monophosphate form of Remdesivir, prime and template were annotated.

**Figure S7.**
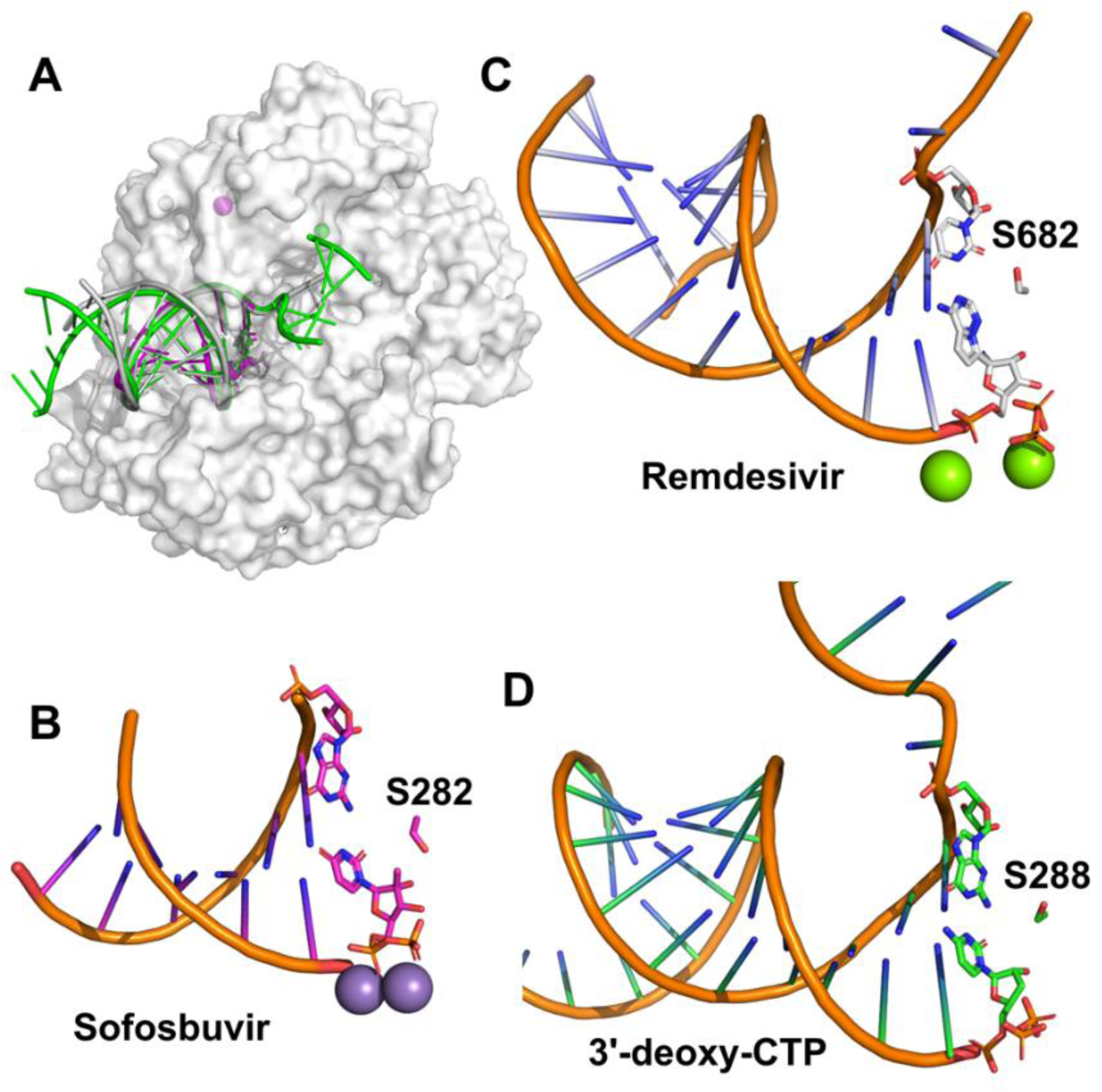
Comparison of SARS-CoV-2 RdRp-RNA complex with HCV RdRp-RNA complex (PDB ID: 4WTF) and poliovirus RdRp-RNA complex (PDB ID: 3OL9). A, A superimposition of three complex structures based on the alignment of RNA. Only the molecular surface of SARS-CoV-2 RdRp is shown. The bound RNA in SARS-CoV-2, HCV, and poliovirus RdRp is colored green, magenta and grey, respectively. B-D, The RNA and the bound ligand in the RdRp-RNA complex of HCV (B), SARS-CoV-2 (C) and poliovirus (D) are shown as cartoon and sticks, respectively. The base paired with the bound ligand and a serine nearby are shown as sticks too.

**Figure S8.**
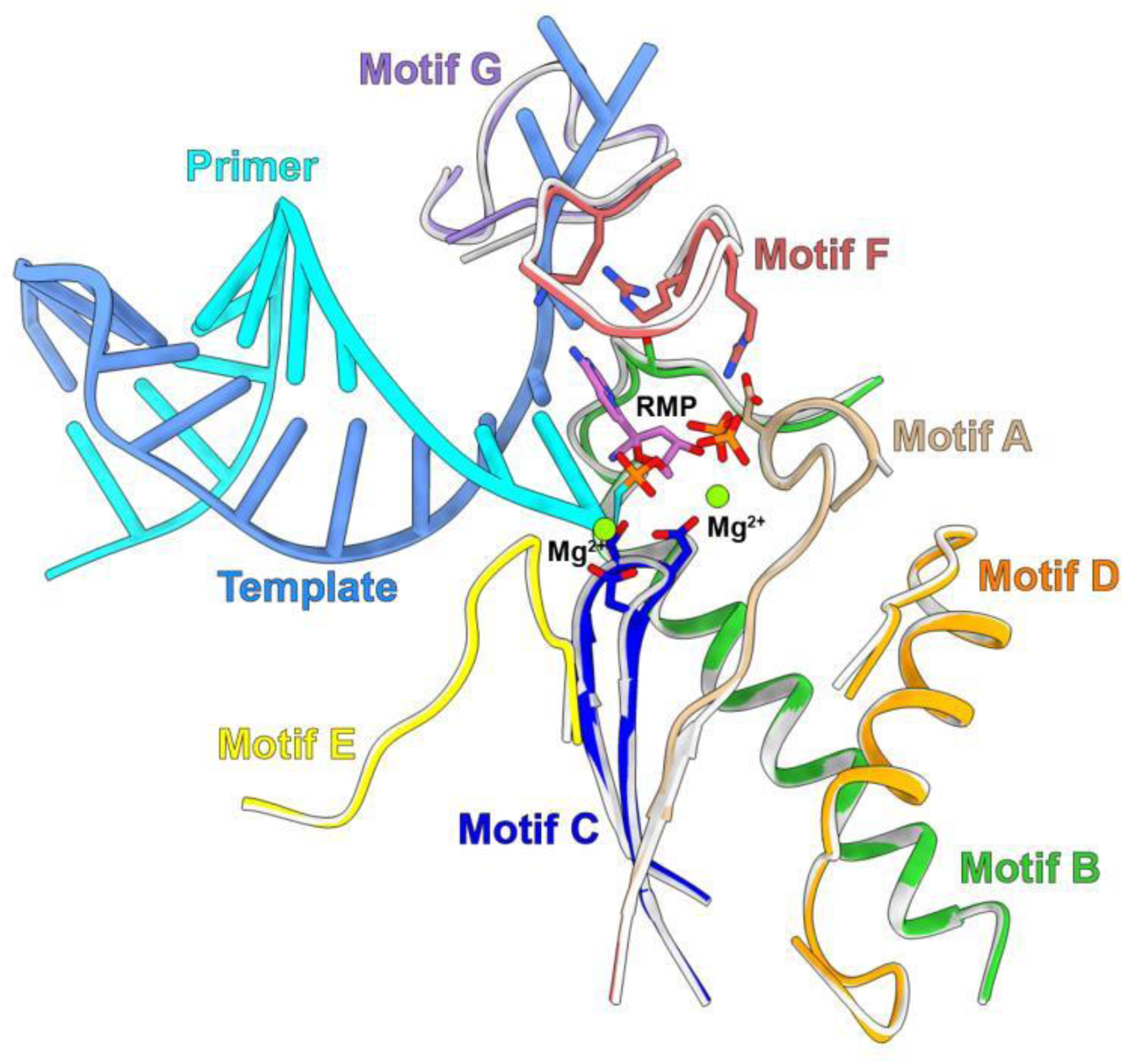
Superimpose of the seven conserved motifs between the apo structure and the RNA bond complex structure. Motifs A to G of the apo RdRp complex are shown as ribbon and colored as grey. Motifs A to G of the RNA bound RdRp complex are colored as tan, green, blue, orange, yellow, dark red and purple, respectively. Remdesivir and key residues of the active site are show in sticks, with the two Mg2+ are shown as sphere and colored as green.

## Supplemental Table 1

**Supplemental Table S1.** EM data and Structure refinement Statistics. **Cryo-EM data collection, model refinement and validation statistics**

| nsp12-nsp7-nsp8 complex | Apo | RNA and RTP bound |
| --- | --- | --- |
| **Data collection and processing** |  |  |
| Magnification | 4,9310 | 4,9310 |
| Voltage (kV) | 300 | 300 |
| Electron exposure (e<sup>-</sup>/Å<sup>2</sup>) | 64 | 64 |
| Defocus range (μm) | -0.8 ~ -3.0 | -0.8 ~ -3.0 |
| Pixel size (Å) | 1.014 | 1.014 |
| Symmetry imposed | C1 | C1 |
| Initial particle projections (no.) | 5,791,192 | 2,438,446 |
| Final particle projections (no.) | 81,494 | 130,386 |
| Map resolution (Å) | 2.8 | 2.5 |
| FSC threshold | 0.143 | 0.143 |
| Map resolution range (Å) | 2.5-5 | 2.2-5 |
| **Refinement** |  |  |
| Initial model used (PDB code) | 6NUR | 6NUR |
| Model resolution (Å) | 3.0 | 2.7 |
| FSC threshold | 0.5 | 0.5 |
| Model resolution range (Å) | 2.6-5 | 2.3-5 |
| Map sharpening *B* factor (Å<sup>2</sup>) | 66.4941 | 57.0431 |
| **Model composition** |  |  |
| Non-hydrogen atoms | 8702 | 8555 |
| Protein residues | 1102 | 1006 |
| Nucleotides | 0 | 25 |
| Water | 0 | 5 |
| Ligands |  |  |
| Zn<sup>2+</sup> | 2 | 2 |
| Mg<sup>2+</sup> | 0 | 3 |
| POP | 0 | 1 |
| RMP | 0 | 1 |
| ***B* factors (Å<sup>2</sup>)** |  |  |
| Protein | 53.26 | 38.60 |
| Nucleotide | --- | 58.70 |
| Ligand | 56.82 | 47.84 |
| Water | --- | 30.00 |
| **R.m.s. deviations** |  |  |
| Bond lengths (Å) | 0.004 | 0.005 |
| Bond angles (°) | 0.675 | 0.658 |
| **Validation** |  |  |
| MolProbity score | 1.69 | 1.50 |
| Clashscore | 6.69 | 5.60 |
| Poor rotamers (%) | 0.00 | 0.00 |
| **Ramachandran plot** |  |  |
| Favored (%) | 95.39 | 96.78 |
| Allowed (%) | 4.61 | 3.22 |
| Disallowed (%) | 0.00 | 0.00 |

